# N-acylethanolamine acid amidase inhibition reduces SARS-CoV-2 infection in Human Precision cut-lung slices and downregulates NF-κB signalling

**DOI:** 10.1101/2024.07.10.602842

**Authors:** Veronica La Rocca, Carolina Filipponi, Viktoria Diesendorf, Alessandro De Carli, Giulia Sciandrone, Silvia Nottoli, Erika Plicanti, Rossella Fonnesu, Elena Iacono, Alessandro Mengozzi, Stefano Masi, Paola Lenzi, Francesco Fornai, Katherina Sewald, Helena Obernolte, Jochen Bodem, Giulia Freer, Mauro Pistello, Michele Lai

## Abstract

Like other positive-sense RNA viruses, SARS-CoV-2 manipulates host lipid metabolism to facilitate its replication by enhancing lipogenesis and lipid droplet formation. In doing so, SARS-CoV-2 infection perturbs bioactive lipid levels associated with the inflammatory response. One of these, Palmitoylethanolamide (PEA) is suppressed during SARS-CoV-2 infection since it activates the Peroxisome Proliferator-Activated Receptor-α (PPAR-α), a transcription factor that suppresses the nuclear factor-B (NF-κB), which is mandatory to sustain SARS-CoV-2 replication. PEA levels are regulated by N-acylethanolamine acid amidase (NAAA), a lysosomal enzyme responsible for catalysing the breakdown of PEA. We hypothesized that NAAA inhibition might interfere with SARS-CoV-2 replication since it will lead PEA to accumulate, activating PPAR-α and, consequently, suppressing NF-κB.

Our results reveal that genetic or chemical ablation of NAAA significantly suppresses SARS-CoV-2 replication by three log_10_ in human-derived precision-cut lung slices. Therefore, we investigated whether inhibiting NAAA could influence NF-κB activation through the activation of PPAR-α. We observed PPAR-α increased expression in NAAA-/-cells, while PPAR-α expression remained low in infected parental cells. As expected, the elevated PPAR-α expression correlated with a parallel reduction in NF-κB activation when NAAA is ablated. These findings underscore NAAA as an essential host factor for SARS-CoV-2 replication and propose a potential mechanism of action rooted in the attenuation of NF-κB activation during viral replication.

**Author summary:** Over the past three years, COVID-19 has claimed nearly 7 million lives worldwide, prompting extensive efforts to find effective treatments. While RNA-based vaccines have been developed rapidly, they alone have not completely halted the spread of the virus, making the search for antiviral therapies crucial. One promising approach targets the anti-inflammatory lipid PEA, which has shown some success in COVID-19 clinical trials. PEA is quickly degraded by the enzyme NAAA. Researchers have found that inhibiting NAAA can enhance and prolong PEA anti-inflammatory effects. NAAA inhibitors have already shown effectiveness in reducing chronic pain and lung inflammation in animal models and have also been effective against Zika virus replication. Our research focused on testing the NAAA inhibitor ARN726 against SARS-CoV-2. In human lung cells and lung tissue samples, ARN726 significantly reduced SARS-CoV-2 replication and inflammation. We discovered that this inhibition suppresses the NF-κB pathway, which the virus uses to fuel its replication and sustain Cytokine storm. Overall, our findings suggest that NAAA inhibitors like ARN726 could be repurposed to combat COVID-19 and potentially other coronaviruses, offering a novel and effective antiviral strategy.

## Introduction

Since the first cases of COVID-19, caused by Severe Acute Respiratory Syndrome Coronavirus 2 (SARS-CoV-2), significant efforts have been made to prevent infection and reduce severe illness through the rapid development of novel or repurposed antiviral drugs. We demonstrated that inhibiting the NAAA enzyme serves as a dual antiviral and anti-inflammatory strategy against Zika virus (Lai et al. 2023). Once NAAA is inhibited, its metabolite PEA, which is no longer hydrolyzed by the enzyme, activates a soluble transcription factor named PPAR-α (Lo Verme et al. 2005). Active PPAR-α causes a shift from lipogenesis to β-oxidation, counteracting viral-induced lipid droplet accumulation. Interestingly, active PPAR-α acts mutually exclusive to NF-κB. Indeed, while NF-κB boosts inflammation, PPAR-α resolve it. Therefore, addressing both viral replication and the hyperinflammatory response typical of COVID-19 could be a valid strategy to counteract Coronavirus diseases effectively.

In Keeping, recent studies uncovered that SARS-CoV-2 ORF3, ORF7, and N proteins specifically activate NF-κB, generating optimal conditions for the virus to replicate and, simultaneously, driving the upsurge of the proinflammatory cascade (Su, Wang, and Yoo 2021; Nilsson-Payant et al. 2021). Moreover, we showed that PEA decreases SARS-CoV-2 infectivity by binding to the S protein in the receptor-binding domain, interfering with its interaction with the ACE-2 receptor (Fonnesu et al. 2022). PEA has been proposed as a dietary supplement to reduce inflammation and SARS-CoV-2 viremia in patients (Fonnesu et al. 2022). Unfortunately, PEA concentrations are quickly reduced by NAAA, which is highly active in immune cells during inflammation.

Several NAAA inhibitors have been developed to overcome this limitation over the years, showing remarkable efficacy in reducing lung inflammation in mice and humans (Wu et al. 2019; Sasso et al. 2018). Since SARS-CoV-2 relies on NF-κB signalling and lipid droplet accumulation, we hypothesized that NAAA inhibitors might exert antiviral activity against SARS-CoV-2 by suppressing NF-κB and activating the disruption of lipid droplets (Dias et al. 2020). Here we show that NAAA inhibition decreases SARS-CoV-2 replication by 4 Log_10_ in human cells *in vitro* and by 3 Log_10_ in human precision-cut lung slices (PCLS), infected *ex vivo*. At the same time, NAAA inhibition reduces the activity of NF-κB by steering PPAR-α activation. These findings pave the way to the possibility of repurposing NAAA inhibitors from drugs with anti-inflammatory properties to potential antiviral activity, especially against single-stranded positive sense (+ss) RNA virus.

## Methods

### Cell culture, treatment, and transfection

Huh-7, Calu-3, and HEK-293T cells were cultured in Dulbecco-Modified Eagle’s Medium (DMEM) supplemented with 10% fetal bovine serum (FBS), 100 U/mL penicillin, 100 μg/mL streptomycin and maintained at 37°C in 5% CO_2_. THP-1 cells were cultured in Roswell Park Memorial Institute 1640 (RPMI-1640, Euroclone, Italy.) Medium supplemented with 10% fetal bovine serum (FBS, Euroclone, Italy), 2mM L-Glutamine, and maintained at 37°C in 5% CO_2_. THP-1 cells were stimulated toward macrophage THP-1 (mTHP-1) using 20 ng/mL PMA for 48 hours. NAAA inhibitor ARN726 (Cayman Chemical, Item NO. 24259, USA) and PPARα inhibitor GW6471 (Cayman Chemical, Item NO. 11697, USA) were diluted in DMSO prior to experiments. To test the toxicity of ARN726, cells were seeded in 96-well plated and treated with ARN726 at 10 μM and 30 μM or DMSO. Cell viability was assessed by MTS assay (CellTiter 96® AQueous One Solution Cell Proliferation Assay (MTS), G3580; Promega; Germany) after 72 h post-treatment. Absorbance was measured at 490 nm, after 1h incubation. Huh-7 and HEK-293T cells were seeded in 6-well plates and transfected with pNL3.2. NF-κB -RE NlucP/ NF-κB-RE/Hygro Vector (N111A Promega, Germany) using Turbofect Transfection Reagent (R0531, ThermoFisher, Germany) according to manufacturer’s instruction.

### Generation of Huh-7 NAAA KO cells

Huh-7 cells were obtained by CRISPR/Ca9 RNP transfection using NEON Transfection system (Thermo Fisher, Massachusetts, USA) using the sgRNA guide: AGTGGGTGCACGTGTTAATC, as described before (Lai et al. 2023).

### Human Precision-Cut Lung slices (PCLS)

PCLS were obtained from 3 donors and infected as previously described (Geiger et al. 2022; Neuhaus et al. 2018). PCLS were kept in a 48-well plate in DEMEM F/12 media (Life Technologies, Germany) supplemented with 1% of penicillin/streptomycin. PCLS were treated with ARN726 at 30 μM and subsequently infected with SARS-CoV-2 at 10^5^ PFU. Media were changed after 24h supplemented with ARN726 at 30 μM. Supernatants from PCLS were collected 72h post-infection. Then, 100 μL were used to infect Vero cells to assess viral production. Viral RNA was extracted after 72h post-infection and quantified by RTqPCR.

### Infections and viral RNA quantification

Huh-7 and Calu-3 cells were seeded in a 48-well plate in a complete medium. The cells were infected with SARS-CoV-2 at 1 Multiplicity of Infection (MOI). 250 μL of supernatants were collected 3 days post-infection. Viral RNA was purified with High Pure Viral Nucleic Acid Kit according to the manufacturer’s instruction (ROCHE, Mannheim, Germany). Viral RNA was amplified by LightMix® Modular Sarbecovirus E-gene (Roche, Mannheim, Germany, Cat. -No. 50-0776-96) using LightCycler 480 II (Roche, Mannheim, Germany).

### RNA extraction and Real-Time PCR

Total RNA was extracted from cell lysate using QIAzol Lysis Reagent (QIAGEN, Hilden, Germany). 200 ng of total RNA was reverse-transcribed in cDNA and then amplified using QuantiNova SYBR Green RT-PCR kit (QIAGEN®, Hilden, Germany). Primers: PPAR-α: F: 5’-CTATCATTTGCTGTGGAGATCG-3’, R: 5’- AAGATATCGTCCGGGTGGTT-3’; hIL8 F:5’-CCTGATTTCTGCAGCTCTGTG-3’, R:5’-CCAGACAGAGCTCTCTTCCAT-3’; NLRP3 F:5’- CTTCTCTGATGAGGCCCAAG-3’, R:5’-GCAGCAAACTGGAAAGGAAG-3’; TNF-α F:5’-AGGAGAAGAGGCTGAGGAACAAG-3’, R:5’-GAGGGAGAGAAGCAACTACAGACC-3’; CXCL10 F:5’- GGAAATCGTGCGTGACATTA-3’, R:5’- AGGAAGGAAGGCTGGAAGAG-3’; IL-6 F:5’-CCAGGAGAAGATTCCAAAGATG-3’, R:5’-GGAAGGTTCAGGTTGTTTTCTG- 3’; IL-10 F:5’-GGGGGTTGAGGTATCAGAGGTAA-3’, R:5’- GCTCCAAGAGAAAGGCATCTACA-3’; TGF-β F:5’-GACATCAAAAGATAACCACTC- 3’, R:5’-TCTATGACAAGTTCAAGCAGA-3’; ACOX1 F:5’- TCCTGCCCACCTTGCTICAC-3’, R:5’-TTGGGGCCGATGTCACCAAC-3’; UCP2 F:5’- CACCAAGGGCTCTGAGCATG-3’, R:5’-TCTACAGGGGAGGCGATGAC-3’; LCAD F:5’- TGAGCGACTGGTGGGAGGAG-3’, R:5’- CACTGTCTGTAGGTGAGCAACTG-3’; β-actin: F: 5’AGGAGAAGCTGTGCTACGTC- 3’, R: 5’-AGACAGCACTGTGTTGGGGTA-3’. Primers and probe: SARS-CoV-2: F:5′- TCACCTAATTTAGCATGGCCTCT-3′,R: 5′-CGTAGTGCAACAGGACTAAGC- 3′,Probe5′-/56FAM/ACAGCAGAATTGGCCCTTAAA/3BHQ-1/-3’. Relative quantification of gene expression was calculated by the 2-ΔΔCt method.

### NF-κB activity: Luciferase assay

Huh-7, Huh-7 NAAA KO cells, and HEK-293T cells were seeded into a 6-well plate (400.000 cells/well). Then, cells were transfected with pNL3.2.NF-κB-RE NlucP/NF- κB-RE/Hygro vector. Cells were then seeded into a 96-well plate, treated or not with the ARN726 30 μM for 30 min before the stimulation with PMA 20 ng/mL. NF-kB Luciferase activity was measured after 24h post-treatment, using Bright-Glo™ Luciferase Assay System (E2610, Promega) according to the manufacturer’s instruction.

### High content confocal imaging

Huh-7 and Calu-3 cells were seeded in 96-well plate 96-CellCarrierUltra plates (Perkin Elmer, Hamburg, Germany), treated with ARN726 30 μM or GW6471 1μM and infected with SARS-CoV-2 1 MOI and 0.1 MOI respectively. Anti-SARS-CoV-2 spike protein (Sino Biological, SI40150-R007-100, Beijing, China, 1:200) or Anti-SARS-CoV-2 Nucleocapsid protein (Sino biological, SI40588-RC02-100, Beijing, China, 1:1000) and the secondary antibodies Alexa488 (Invitrogen, A11008, 1:1000) or Alexa647 (Invitrogen, A21245, 1:1000) were used to stain SARS-CoV-2. LDs were stained using OIL Red R (1:5000) for 15 min and then washed in water. Nuclei were stained with DAPI (1 μg/mL). Images were acquired using Operetta CLS high-content imaging device (PerkinElmer, Hamburg, Germany) and analyzed with Harmony 4.6 software (PerkinElmer Hamburg, Germany) accordingly to the following building block: Find Nuclei > Find cytoplasm > Calculate intensity properties (AlexaFluor488/647) > Select population AlexaFluor488/647 (Infected cells) > Find Spot (LD-OIL Red) > Calculate position properties (% overlap infected cells-OIL Red).

### Transmission electron microscopy (TEM)

Huh-7 cells were infected with SARS-CoV-2 as described above. Then, cells were fixed with 2.0% paraformaldehyde/0.1% glutaraldehyde, both dissolved in 0.1 M PBS pH 7.4 for 90 min at 4 °C. Cells were gently scraped from the plate, collected into vials, and centrifuged at 10,000 rpm for 10 min. Cells were then resuspended in PBS and post-fixed in 1% osmium tetroxide (OsO_4_) for 1h at 4 °C. Then, the post-fixed pellet was dehydrated in a gradient of ethanol solutions of 50 %, 70 %, 90 %, and 95 %, each for 5 min, to reach 100% ethanol for 60 min. Finally, the pellet was embedded in epon araldite for 72 h, at 60°C. The embedded cell samples were sectioned at the ultramicrotome (Leica Microsystems) and ultrathin sections underwent conventional electron microscopy or immune electron microscopy. Samples were observed at Jeol JEM SX100 TEM (JEOL, Tokyo, Japan) at an acceleration voltage of 80 kV.

### Statistical analysis

Statistical significance of differences between the groups was performed using Student’s t-test or One-Way ANOVA, followed by Bonferroni post-test (multiple comparisons) using Graph-Pad PRISM 9 software. Differences between groups were considered statistically significant at p values < 0.05. Results are expressed as mean ± SD of two or three independent experiments.

## Results

### NAAA ablation reduces SARS-CoV-2 replication

To investigate the role of NAAA in SARS-CoV-2 replication, we used Huh-7 as they naturally support SARS-CoV-2 replication (Zhou et al. 2020). Subsequently, we infected both Huh-7 NAAA knockout (KO) and the wild type (WT) cells with SARS-CoV-2 at 1 MOI and quantified viral genomes in the supernatants using RT-qPCR. We found a 5-fold reduction in SARS-CoV-2 genome copy content in Huh-7 NAAA KO cells compared to their WT counterparts (Figure 1b). Furthermore, we observed that the SARS-CoV-2 nucleocapsid (N) protein amounts in cell lysates exhibited a decrease of approximately 1 Log_10_ in Huh-7 NAAA KO cells compared to the WT cells (Figure 1c-d). Thus, to shed light on how NAAA activity might interfere with SARS-CoV-2 replication, we analyzed Huh-7 NAAA KO cells and their WT counterparts by TEM. Huh-7 NAAA KO cells accumulated a higher number of vesicle-shaped organelles in their cytoplasm, suggesting an accumulation of autophagosomes/autolysosomes or peroxisomes (Figure 1e-h).

**Figure 1.**
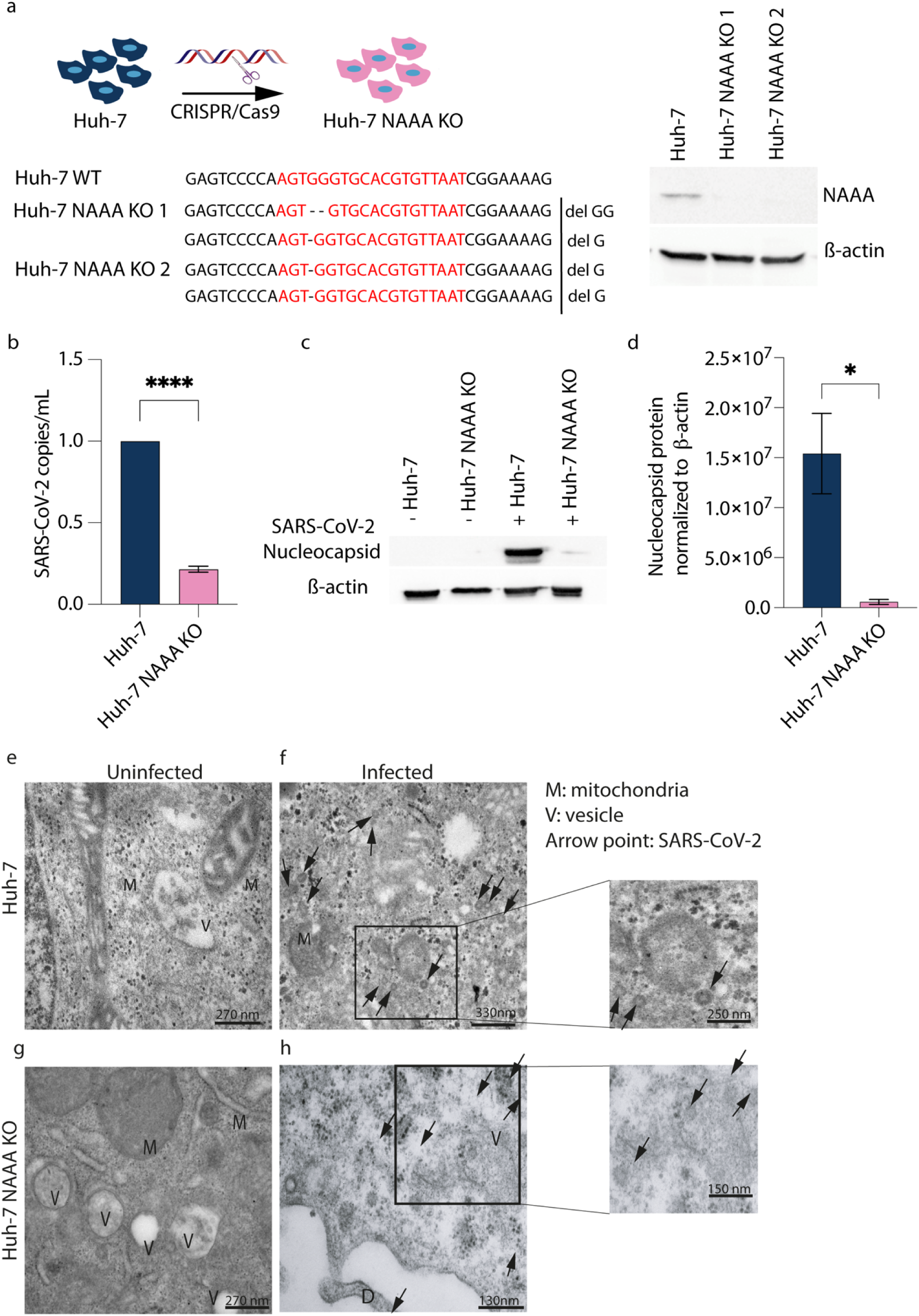
NAAA ablation reduced SARS-CoV-2 infection. a) Left: Schematic illustration of deletions in Huh-7 NAAA KO cells obtained by CRISPR-Cas9 transfection. Red writing indicates the target gRNA sequence on gene ASAHL, encoding NAAA. Hyphens represent nucleotide deletions. Right: Western blot analysis shows the effects of ASAHL genetic disruption at protein level in Huh.7 NAAA KO cells scoring negative for NAAA. b) Huh-7 NAAA KO cells and their WT counterparts were infected or not with SARS-CoV-2, then, at 48h post-infection, RT-qPCR was performed on supernatants to quantify SARS-CoV-2 genomes. c) Western blot analysis was performed against SARS-CoV-2 N protein in cell lysates. d) Analysis of western blot in c). Results are expressed as mean ± SD of 3 independent replicates. Data were analysed by Student’s T-test (*p<0.0.5; ****p<0.0001). e-f) Representative TEM images of e-g) uninfected cells and f-h) SARS-CoV-2 infected cells analyzed 48 hpi. Scale bar: 270nm.

### NAAA inhibitor ARN726 decreases SARS-CoV-2 replication in cell culture

To assess whether NAAA inhibition might affect SARS-CoV-2 replication, we pretreated Calu-3 cells with a potent inhibitor of NAAA (ARN726) prior to infection. These cells are commonly used to mimic SARS-CoV-2 infection of human lung cells. Cells were then treated with ARN726 and GW6471, a PPAR-α inhibitor that might counteract the activation of pathways downstream PEA accumulation. Then, cells were infected with SARS-CoV-2 and analyzed by High-Content Confocal screening (Figure 2a). First, we confirmed the lack of cytotoxicity of ARN726 and GW6471 on Calu-3 cells, as previously reported (Figure 2b) (Lai et al. 2023). The analysis revealed that NAAA inhibition decreased the number of infected cells by ∼1 Log_10_, while GW6471 did not exert any antiviral activity (Figure 2c-e). Surprisingly, co-administration of ARN726 and GW6471 decreased SARS-CoV-2 replication as much as ARN726 alone, indicating that PPAR-α activation might not be the only mechanism against SARS-CoV-2 replication acting downstream NAAA inhibition in our model. Finally, we measured the copies of viral genomes released in the supernatants 72 hpi. As shown in Figure 2e, we found a 4 Log_10_ reduction in SARS-CoV-2 replication in Calu-3 cells, compared to the DMSO-treated counterparts.

**Figure 2.**
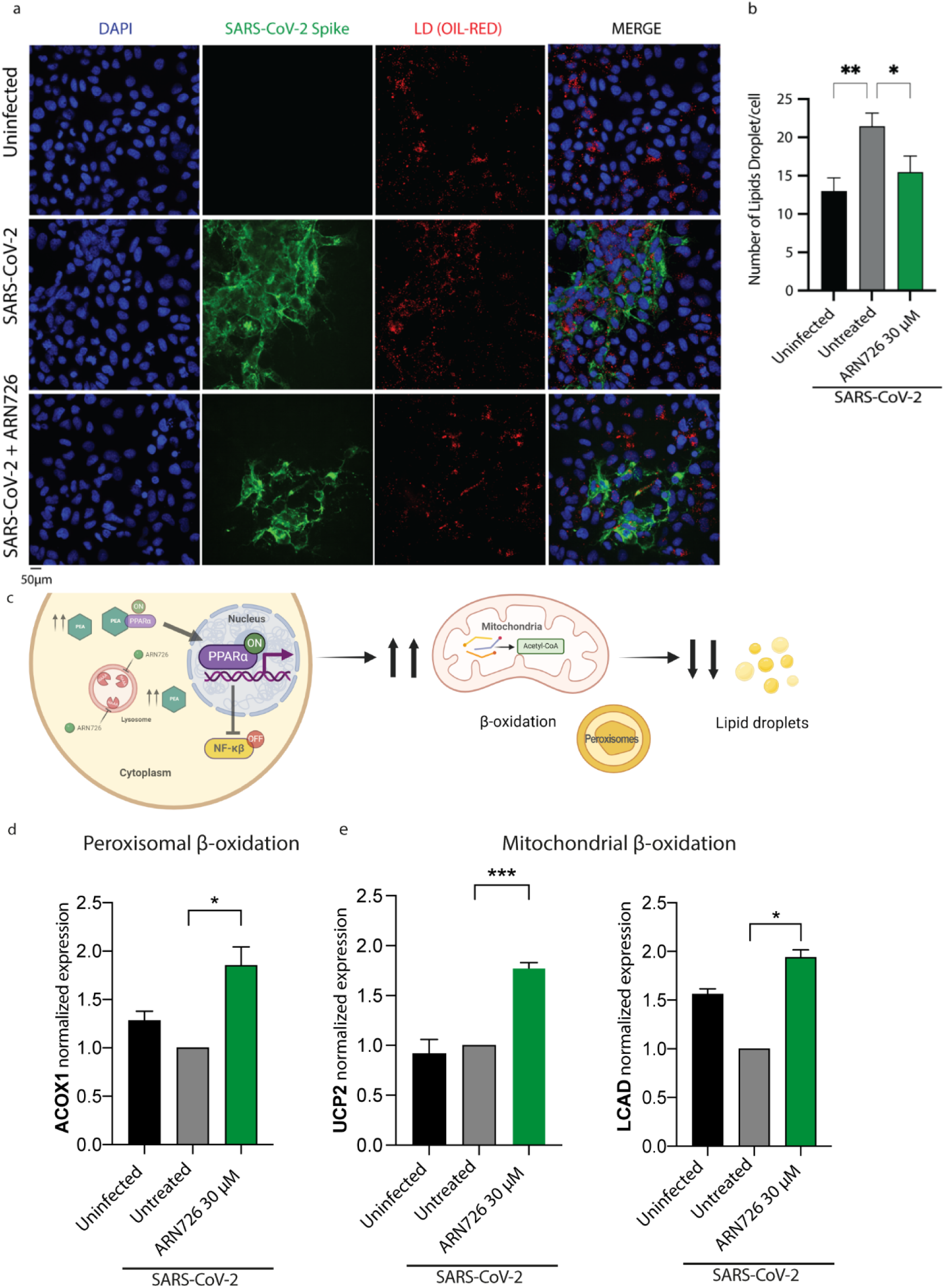
ARN726 reduces SARS-CoV-2 replication. a) Illustration of the experimental workflow: Calu-3 cells were treated with ARN726 at 30 μM for 30 min and then infected with SARS-CoV-2. Supernatants were collected 72h post-infection and analyzed by RT-qPCR. b) Calu-3 cells were treated with ARN726 at 10 and 30 μM, GW6471 at 1, 5, 10, 50, and 100 μM or only the vehicle (DMSO) for 72h. Cell viability was analysed by WST assay measuring the absorbance at 490 nm. c) Representative images of Calu-3 cells stained for SARS-CoV-2 N protein (Alexa Fluor-647), phalloidin (Alexa Fluor-488), and nuclei (DAPI). d) Statistical analysis of SARS-CoV-2 positive cells /total cell number. Images were acquired using a 40x water objective. e) Calu-3 cells were treated or not with ARN726 30μM and infected with SARS-CoV-2. Viral loads were determined in supernatants 72 hpi by RT-qPCR. Results are expressed as mean ± SD of independent replicates. Data were analysed with Student’s t-test or One-Way ANOVA (***p<0.001; ****p<0.0001).

### ARN726 reduces SARS-CoV-2 replication in Human Precision Cut Lung Slices (PCLS)

To further validate the antiviral activity of ARN726 against SARS-CoV-2, we tested the compound on PCLS, the closest 3D model to human lung structure, maintaining its specific multi-cellular composition. PCLS were obtained from 3 uninfected human donors as previously described (Zimniak et al. 2021; Pechous et al. 2023). As shown in figure 3a, PCLS were treated with ARN726 at 30 μM and first tested for cytotoxicity using MTS assay (Figure 3b). Then, PCLS were infected with SARS-CoV-2 at 0.1 PFU/mL. Afterward, viral load was determined on Vero cells (Figure 3c). We found a 3 Log_10_ reduction in SARS-CoV-2 viral load in PCLS treated with ARN726 at 30 μM compared to the untreated controls.

**Figure 3.**
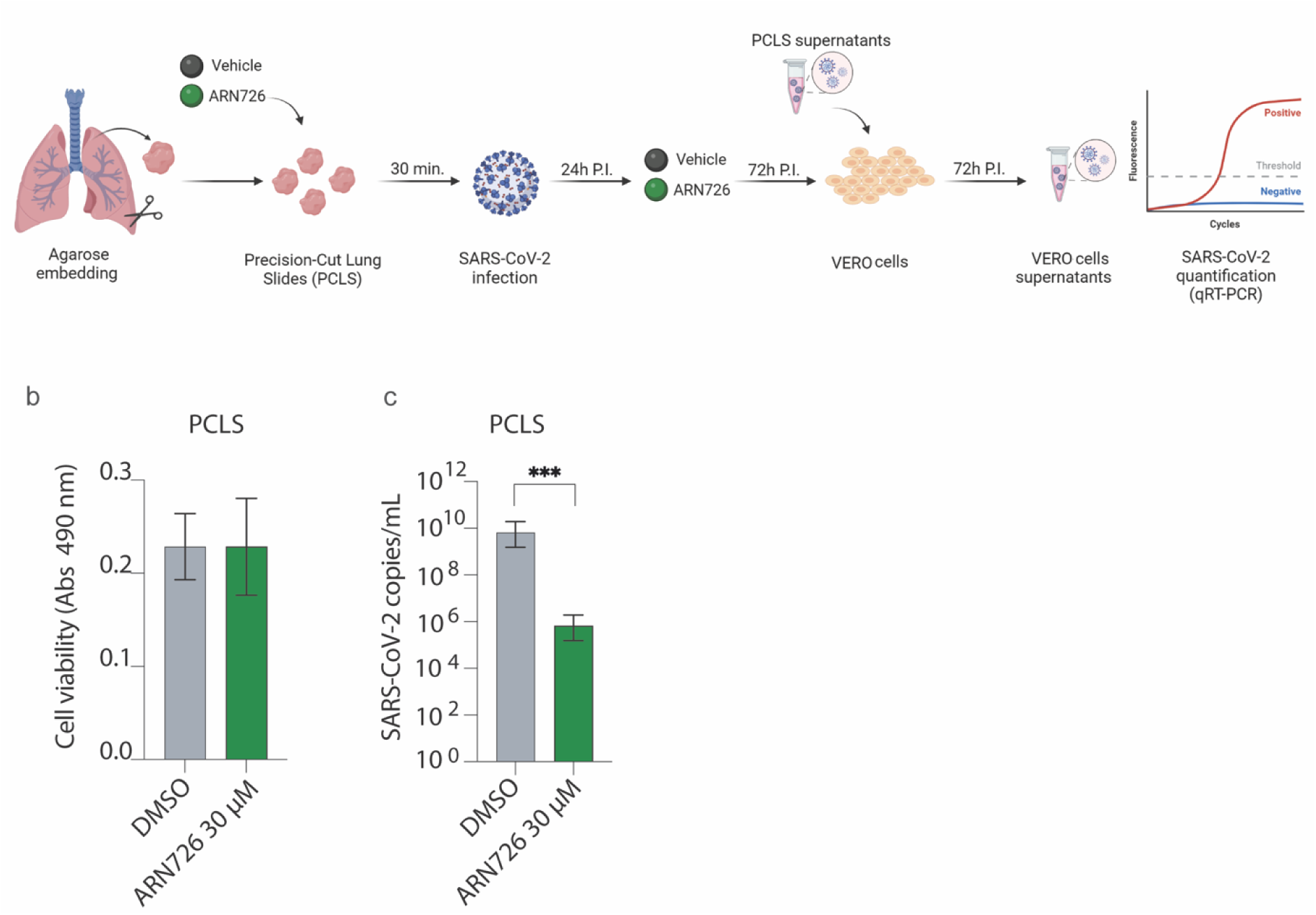
ARN726 reduces SARS-CoV-2 replication in PCLS. a) Schematic illustration of the experimental workflow. Briefly, PCLS were treated with ARN726 30 μM or DMSO and infected with SARS-CoV-2. Media with or without compound were changed after 24 hpi. Then 100 μL of supernatants were collected and used to infect VERO E6 cells. Viral RNA was extracted from supernatants and amplified by RT-qPCR. b) viability was assessed, administering ARN726 at 30 μM for 3 days. c) Viral load was assessed by RT-qPCR 72 hpi. Results are expressed as the mean ± SD of 3 independent replicates. Data were analysed with a student’s t-test (***p<0.001).

### NAAA inhibition downregulates NF-κB activation by steering PPAR-α pathway

It has been recently reported that SARS-CoV-2 ORF3, ORF7, and N proteins activate NF-κB (Su, Wang, and Yoo 2021). NF-κB activation causes a cascade of events that ultimately contribute to the excessive production of pro-inflammatory chemokines and cytokines, leading to the severe outcome of COVID-19 (Nilsson-Payant et al. 2021). In keeping, several NF-κB inhibitors exhibit remarkable antiviral activity against SARS-CoV-2 (Gudowska-Sawczuk and Mroczko 2022). However, the complex biology of NF-κB, a key regulator of many cellular processes involved in cell survival and proliferation, makes it a questionable pathway to for therapeutic purposes, a fact confirmed by the lack of FDA-approved drugs acting on NF-κB (Nilsson-Payant et al. 2021).

It is widely recognized that NAAA inhibitors effectively downregulate NF-κB activation by modulating PPAR-α, which governs a mutually exclusive pathway to NF-κB (Lo Verme et al. 2005). Indeed, PEA-activated PPAR-α suppresses NF-κB transcription. Based on this evidence, we hypothesized that the antiviral activity observed for NAAA inhibitors might depend on the suppression of NF-κB -related pathways that indirectly affect SARS-CoV-2 replication, as schematically reported in Figure 4a. Thus, we measured PPAR-α gene transcription in Huh-7 NAAA KO cells and their WT counterparts before and after SARS-CoV-2 infection. Infection with SARS-CoV-2 Wuhan and B.1.1.7 variants suppresses PPAR-α activation in WT cells whereas it is enhanced in NAAA KO cells (Figure 4b).

**Figure 4.**
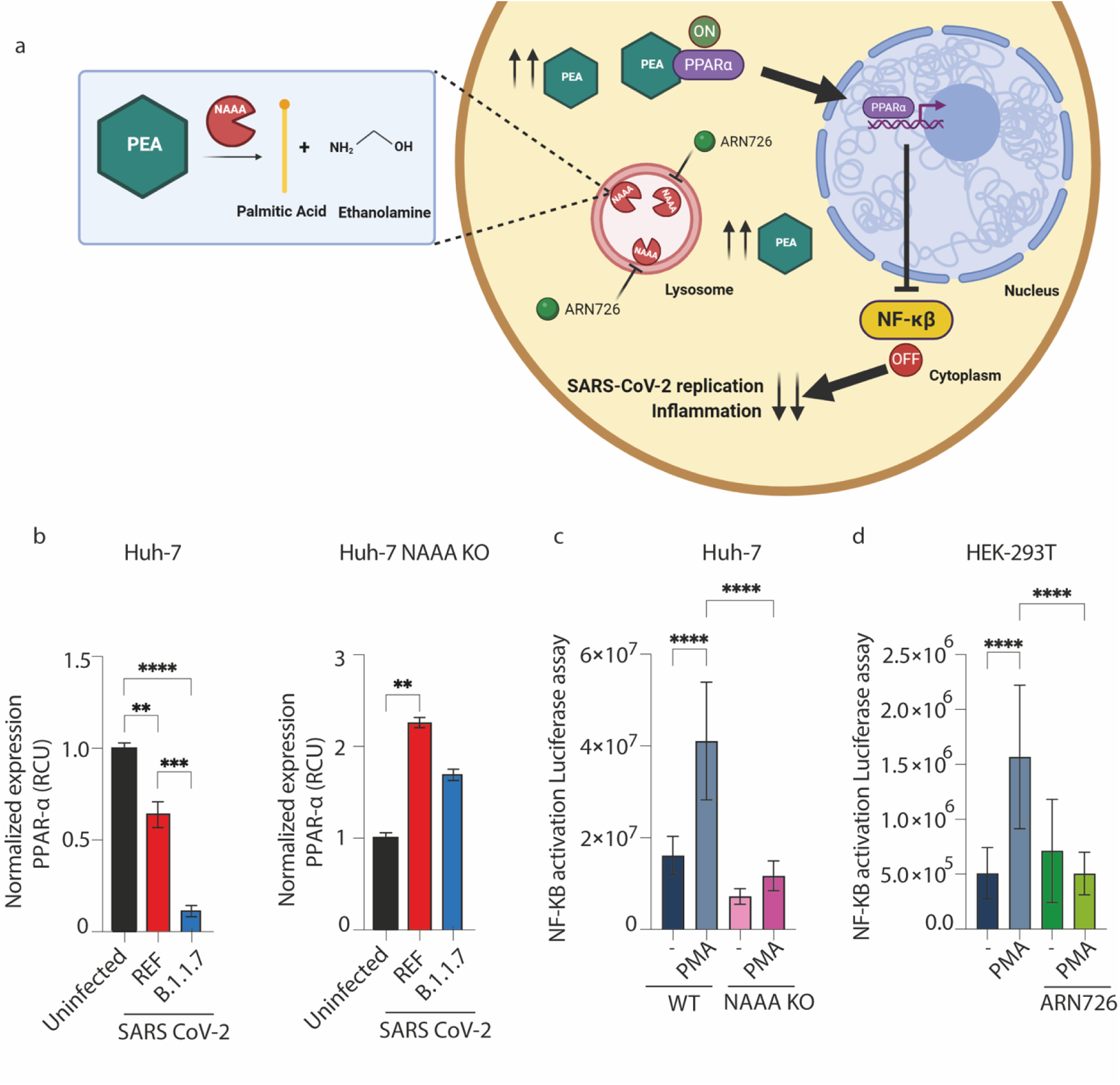
NAAA inhibition increases PPAR-α expression and inhibits NF-κB activation. a) Schematic illustration of the signalling pathway downstream NAAA inhibition: ARN726 inhibits NAAA, raising PEA levels that activate PPAR-α, which in turn suppresses NF-κB. b) Huh7 and Huh-7 NAAA KO cells were infected with SARS-CoV-2 at 1 MOI (Ref: UK strain, and B.1.1.7 strain). PPAR-α mRNA was determined by RT-qPCR and normalized to the uninfected cells using ACTIN as a housekeeping gene. Results are expressed as mean ± SD of three independent replicates. Data were analysed with One-Way ANOVA (** p<0.01; ***p<0.001). (c, d): Huh-7 or HEK-293T cells were transfected with a NF-κB luciferase reporter vector. Then, cells were treated with ARN276 at 30 μM following stimulation with PMA at 20 ng/mL 24 hpi. To measure NF-κB activation luciferase assay was performed. Results are expressed as the mean ±SD of three independent replicates. Data were analysed with One-Way ANOVA (****p<0.0001).

Since PPAR-α suppresses NF-κB activation, we tested whether PPAR-α upregulation in NAAA KO cells was accompanied by NF-κB activity suppression in a luciferase-reporter-based assay. Briefly, we transfected Huh-7, Huh-7 NAAA KO cells, or HEK-293T cells with a plasmid containing five NF-κB responsive elements driving luciferase transcription. Then, cells were treated or not with ARN726 or Phorbol myristate acetate (PMA), a potent NF-κB activator (Zhang et al. 2023). PMA administration doubled NF-κB activation in Huh-7, while it did not exert any effect in NAAA KO counterparts (Figure 4c). This evidence was also confirmed by the administration of ARN726 to HEK-293T cells (Figure4d) exposed to PMA, in this case, NF-κB activation was suppressed when the compound was administered at 30 μM.

### NAAA inhibition decreases LD formation and activates mitochondrial and peroxisomal β-oxidation

It has been demonstrated that interfering with NF-κB signalling results in a dramatic decrease in SARS-CoV-2 replication (Nilsson-Payant et al. 2021). NF-κB has been shown to be the dominant driver of the host transcriptional response in SARS-CoV-2-infected cells and a likely contributor to the underlying inflammation observed in COVID-19 (Dias et al. 2020; Nilsson-Payant et al. 2021). In contrast to PPAR-α, NF-κB activation leads to lipogenesis, causing enhanced LD accumulation and consequent pro-inflammatory eicosanoid release (Pereira-Dutra et al. 2019; Bozza et al. 2011). We hypothesized that the mechanism behind the antiviral activity of NAAA inhibitors relies on the suppression of NF-κB. This may be induced by the activation of the PPAR-α pathway and the consequent disruption of a pro-viral environment, represented by an abundance of LDs, essential for SARS-CoV-2 replication (Wang et al. 2023). To probe so, we treated or not Huh-7 cells with ARN726 at 30 μM. Then, we infected cells with SARS-CoV-2 at 1 MOI, then stained them for SARS-CoV-2 S protein and Oil RED (LDs) and compared the LDs by high-content confocal screening to uninfected cells. This assay confirmed that SARS-CoV-2 doubled LD content in Huh-7-infected cells, while the administration of NAAA inhibitor ARN726 counteracted SARS-CoV-2-induced LD accumulation, maintaining a similar LD quantity and distribution to uninfected cells (Figure 5a-b). To better characterize LD biogenesis in ARN726-treated cells, we quantified the expression of β-oxidation key enzymes ACOX1, UCP2, and LCAD by RT-qPCR (Figure 5c), regulating peroxisomal (ACOX1) or mitochondrial (UCP2 and LCAD) β-oxidation (Figure 5d-e). We found that ACOX1, UCP2, and LCAD mRNAs resulted in up-regulated by ∼ 2-folds.

**Figure 5.**
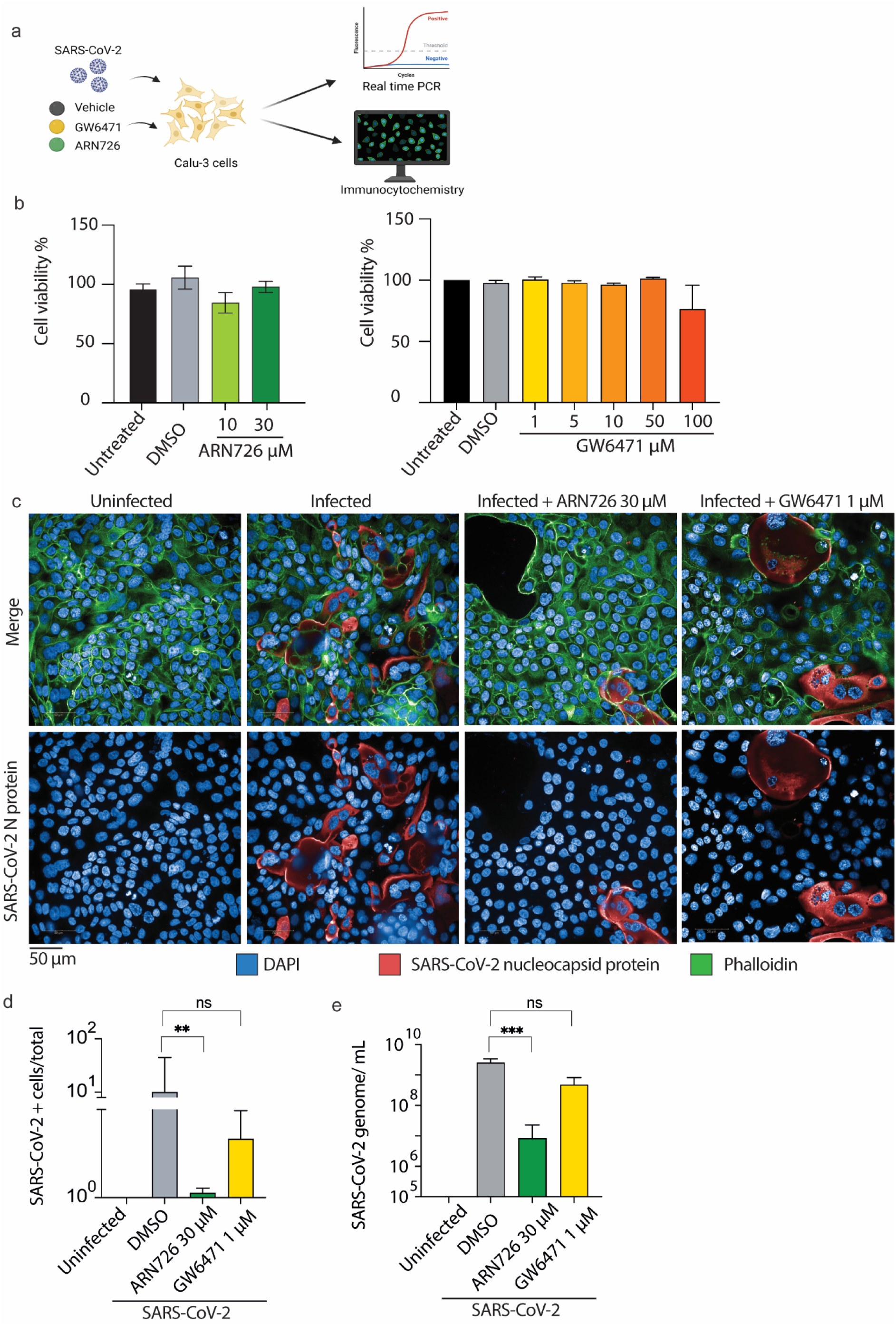
NAAA ablation prevents LD accumulation during SARS-CoV-2 infection. a) representative images of Huh-7 cells stained for SARS-CoV-2 S protein (Alexa Fluor-488), LDs (OIL RED), and nuclei (DAPI). b) Statistical analysis of LD number /cell. The scale bar is depicted below the panel. Data are expressed as mean ± SD of three experiments and analysed by One Way-ANOVA * p<0.05 ** p<0.01. c) Schematic illustration of the mechanism proposed. Briefly, ARN726 blocks NAAA in the lysosomes, accumulating PEA, and activating the transcription factor PPAR-α. As a result, NF-κB expression is suppressed and LDs are used up by β-oxidation. d) quantification analysis of ACOX1, related to peroxisomal β-oxidation by RT-qPCR. e) Quantification analysis of UCP2 and LCAD engaged in mitochondrial β-oxidation by RT-qPCR. Data are expressed as mean ±SD (N=3) and analyzed by One-Way ANOVA (* p<0.05; *** p<0.001).

### NAAA inhibition prevents SARS-CoV-2 cytokine storm

Overactivation of CD4+ and CD8+ T cells and the resulting cytokine and chemokine production lead to cytokine storm, an uncontrolled hyperinflammatory reaction, causing local and distant tissue damage. This heightened inflammation is associated with peripheral blood lymphopenia, a significant drop in the lymphocyte-to-neutrophil ratio, and CD4+ T-cell dysfunction (Copaescu et al. 2020). Inflammatory cytokines and chemokines dysregulated by SARS-CoV-2 infection include IL-6, IL-10, TNF-α, TGF-β, NLRP3, and CXCL10, which were found significantly elevated in patients with severe SARS-CoV-2 infection (Coperchini, Chiovato, and Rotondi 2021). To demonstrate that NAAA inhibition might counteract the SARS-CoV-2-induced cytokine storm, we analysed the mRNA expression of NLRP3, IL-6, IL-10, TNF-α, TGF-β, and CXCL10 in mTHP-1 cells infected with SARS-CoV-2 and then treated or not with ARN726. Overall NAAA inhibition suppresses SARS-CoV-2 induced cytokine storm. Indeed, NLRP3 transcription decreases by 8-folds when ARN726 is administered. Also IL-10, IL-16, TNF-α, and TGF-β, key players of inflammation, remained suppressed in SARS-CoV-2-infected cells treated with ARN726 (Figure7 c-g).

**Figure 7.**
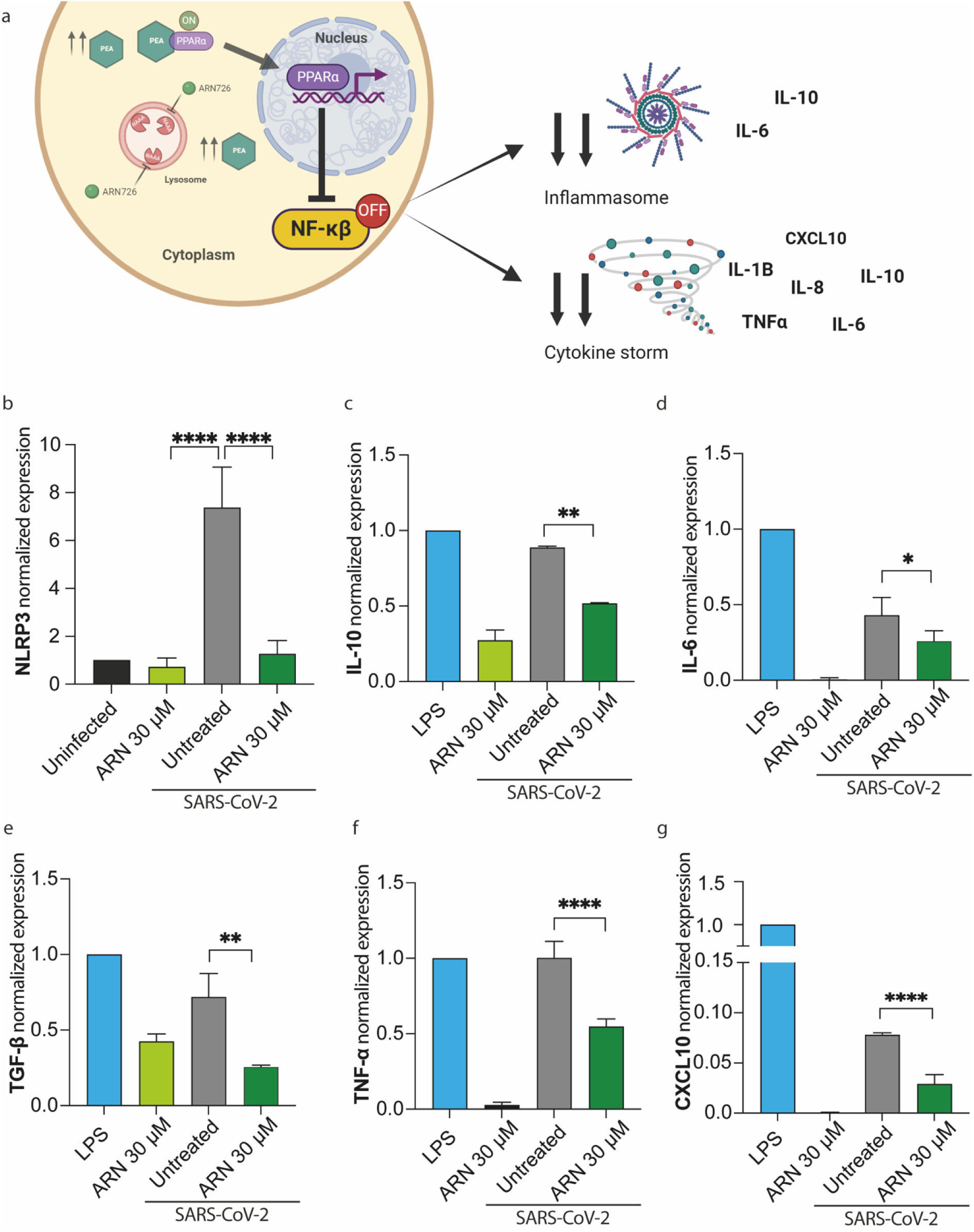
ARN726 reduces SARS-CoV-2 cytokine storm. a) Schematic illustration of the mechanism. Briefly, ARN726 blocks NAAA activity in the lysosomes, leading to PEA accumulation, and activating the transcription factor PPAR-α. As a result, NF-κB expression is suppressed and inflammation-related genes are not expressed downstream. b-g) Expression analyses of NLRP3, IL-10, IL-6, TGF-β, TNF-α, and CXCL10 by RT-qPCR. Data are expressed as mean ±SD (N=3) and analyzed by One-Way ANOVA (* p<0.05; ** p<0.01, *** p<0.001 **** p<0.0001).

## Discussion

Over the past 3 years, COVID-19 has evolved into a global pandemic, resulting in nearly 7 million fatalities worldwide. Throughout the COVID-19 pandemic, substantial efforts have been invested in devising effective strategies against infection, leading to the rapid development of RNA-based vaccines. Furthermore, significant attention has been devoted to drug repurposing strategies, with the goal of pinpointing promising candidate drugs for clinical use. However, it has become increasingly evident, although severe cases were significantly reduced, that the COVID-19 spread has not been solved by vaccination alone since the disease persists in the population. As a result, the need to identify effective antiviral therapies has remained imperative, and significant attention is being devoted to drug repurposing strategies. Several researchers highlighted the use of the anti-inflammatory lipid PEA to mitigate the severe symptoms of COVID-19. PEA is a well-documented endogenous bioactive lipid with anti-inflammatory properties and involvement in various cellular pathways, including those hijacked by viral infections. These established properties have drawn interest from researchers, with PEA having been tested in 2 clinical trials with partial success (Di Stadio et al. 2022). Since PEA is rapidly degraded by NAAA, several independent research groups showed that inhibiting NAAA exerts a stronger and longer-lasting effect on controlling inflammation if compared to the administration of PEA. Consequently, NAAA inhibitors have been developed, and their effectiveness has been demonstrated in alleviating chronic pain, lung inflammation, and neurogenic pain in mouse and rat models. In addition, we have previously demonstrated that NAAA inhibition reduced ZIKV replication and impeded its maturation in neuronal stem cells (Lai et al. 2023). This motivated our investigation into the role of NAAA during SARS-CoV-2 infection. Indeed, like ZIKV, SARS-CoV-2 replication relies on suppressing PPAR-α by steering its opposite pathway, NF-κB. We tested NAAA inhibitor ARN726 against SARS-CoV-2 and found that it significantly reduced viral replication by 4 Log_10_ in Calu-3 cells. Moreover, we further tested ARN726 in PCLS, infected or not with SARS-CoV-2. PCLS were obtained from human lungs and represented a valuable *ex vivo* model to study how SARS-CoV-2 reacts to NAAA inhibition in a context closer to *in vivo* conditions (Pechous et al. 2023). Indeed, PCLS maintain the architecture found in human lungs as closely as achievable *ex vivo*. We observed that NAAA inhibition reduced SARS-CoV-2 replication by 3 Log_10_ in PCLS, confirming the striking antiviral potency of NAAA inhibition observed in both Huh-7 and Calu-3 cells.

To shed light on the mechanism involved in the antiviral activity of NAAA inhibition, we investigated whether a reduction in NF-κB signalling might be involved. This pathway is crucial in SARS-CoV-2 replication since the virus encodes at least 3 proteins devolved to NF-κB activation. Therefore, it is plausible that this pathway might be turned off by NAAA inhibition. Indeed, in NAAA -/-cells, the activation of the NF-κB transcription factor is downregulated even in the absence of any stimulus, whereas NF-κB activation drops soon after NAAA inhibition in HEK293T cells stimulated with PMA.

Once established that the NF-κB pathway is suppressed by NAAA inhibition, we attempted to shed light on cellular events following SARS-CoV-2 infection. Indeed, SARS-CoV-2 replication brings along inflammation and, most importantly for the virus, LD synthesis as fuel and as a source of lipid material for its replication organelle building. Of note, SARS-CoV-2 cytokine storm is also caused by LDs accumulation, since LDs play a critical role in regulating the production of inflammatory factors (Pereira-Dutra et al. 2019). Indeed, it is possible to speculate that NF-κB activation triggered by SARS-CoV-2 boosts lipogenesis, which benefits the virus by fueling its replication. In contrast, when NAAA is inhibited, LDs are destroyed by the activation of PPAR-α, which steers β-oxidation and suppresses NF-κB transcription. Our findings reveal that the absence of NAAA activity led to a remarkable 30% reduction in the number of LDs in SARS-CoV-2-infected cells. Our results align with previous research. (Dias et al. 2020). It is very likely that the reduction in LDs induced by ARN726 during SARS-CoV-2 infection exerts an anti-inflammatory effect. Indeed, we found a striking anti-inflammatory activity in THP-1 cells treated with NAAA inhibitors after SARS-CoV-2 infection. Notably, the mRNA levels of NLRP3, IL-6, IL-10, TGF-β, TNF-α, and CXCL10 did not differ from those in uninfected counterparts.

All together, these findings show that NAAA inhibition has a beneficial effect by both reducing inflammation and blocking viral replication of SARS-CoV-2. Therefore, our research suggests that repurposing NAAA inhibitors could offer a novel approach against new *Coronaviridae* with pandemic potential.

### Institutional Review Board Statement

Human lung lobes were acquired from patients under-going lobe resection for cancer at Hannover Medical School. The use of the tissue for research was approved by the ethics committee of the Hannover Medical School and complies with the Code of Ethics of the World Medical Association (number 2701–2015).

### Informed Consent Statement

All patients gave written informed consent to use explanted lung tissue for research and publish the results. No sample tissues were procured from prisoners.

